# Rapid Prototyping of Thermoplastic Microfluidic 3D Cell Culture Devices by Creating Regional Hydrophilicity Discrepancy

**DOI:** 10.1101/2023.06.23.546122

**Authors:** Haiqing Bai, Kristen N. Peters Olson, Ming Pan, Thomas Marshall, Hardeep Singh, Jingzhe Ma, Paige Gilbride, Yu-Chieh Yuan, Longlong Si, Sushila Maharjan, Di Huang, Xiaohua Qian, Carol Livermore, Yu Shrike Zhang, Xin Xie

## Abstract

Microfluidic three-dimensional cell culture devices that enable the recapitulation of key aspects of organ structures and functions *in vivo* represent a promising preclinical platform to improve translational success during drug discovery. Essential to these engineered devices is the spatial patterning of cells from different tissue types within a confined microenvironment. Traditional fabrication strategies lack the scalability, cost-effectiveness, and rapid prototyping capabilities required for industrial applications, especially for processes involving thermoplastic materials. Here, we introduce an approach to pattern fluid guides inside microchannels by establishing differential hydrophilicity using pressure-sensitive adhesives as masks and a subsequent selective coating with a biocompatible polymer. We identified optimal coating conditions using polyvinylpyrrolidone, which resulted in rapid and consistent hydrogel flow in both the open-chip prototype and the fully bonded device containing additional features for medium perfusion. We tested the suitability of our device for dynamic 3D cell culture by growing human hepatocytes in the device under controlled fluid flow for a 14-day period. Additionally, we demonstrated the potential of using our device for pharmaceutical high-throughput screening applications, such as predicting drug-induced liver injury. Our approach offers a facile strategy of rapid prototyping thermoplastic microfluidic organ chips with varying geometries, microstructures, and substrate materials.

## Introduction

Despite the significant investment in research and development over the past decade, the annual approval rate for new drugs has remained largely static (*1*). A major contributor to the impasse is the low rate of successful translation from preclinical studies to clinical development. Human organ chips as an alternative to traditional animal models commonly used in modern drug development, may offer a solution to this translational challenge by serving as a more faithful preclinical platform to study human disease processes as well as to evaluate drug safety and efficacy (*2, 3*).

Organ chips (or Organ-on-Chips, OOC) are bioengineered devices containing living cells that mimic the structures and core functions of specific organs *in vivo* (*2*). Emerging from advancements in microfabrication, mechanobiology, tissue engineering, and stem cell biology, OOC were initially developed as research tools to model and study disease processes *in vitro*. However, in recent years, they have gained increasing traction in the fields of preclinical disease modeling, drug testing, and personalized medicine. One promising application of OOC in the drug development process is predicting drug-induced liver injury (DILI)-a type of liver injury caused by various drugs, herbs, or other xenobiotics (*4*). DILI is responsible for over 20% of all drug failures, yet current preclinical models, based largely on animal testing, often fail to predict DILI in humans (*5*). In contrast, recent benchmark studies involving 27 known hepatotoxic and non-toxic drugs found that OOC can predict DILI with 100% sensitivity and 87% specificity (*6*). However, a greater adoption of OOC for preclinical DILI testing necessitates the development of more cost-effective and high-throughput OOC devices (*7*).

Current OOC fabrication largely employs the silicone elastomer polydimethylsiloxane (PDMS) (*8*). While PDMS allows for rapid prototyping and has been widely used in the academic setting, it lacks the accessibility and scalability required for large-scale drug testing, and it has issues related to adsorption or absorption of hydrophobic molecules (*9*). Thermoplastic materials, such as polystyrene (PS), polycarbonate (PC), and cyclic olefin copolymer (COC), are better suited due to their low cost, ease of fabrication, and biocompatibility (*8, 10*). However, prototyping microfluidic devices with these materials typically involves micro-molding or injection-molding, which can be expensive and time-consuming. Thus, a fast-prototyping strategy for OOC devices using thermoplastic materials is urgently needed.

A distinctive feature of OOC is the capacity to culture cells from various tissue types in a spatially-controlled manner. This facilitates the replication of complex human organ architectures, as well as the establishment of biochemical gradients, vascularization, and fluid sampling from different compartments. One method to achieve this involves controlling hydrogel flows within OOC microchannels, which eliminates the need for a porous membrane that may hinder cell migration and is not ideal for modeling diseases such as fibrosis, where the composition and physical properties of the extracellular matrix (ECM) change with disease progression (*11*). For example, micropillars (*12*) and phaseguides (*13*) are the two common features that can be integrated into the microchannels to achieve guided hydrogel flow. They work by creating a capillary pressure barrier between compartments, thus allowing one compartment to be filled without the liquid spilling over into the adjacent compartment. However, these features also have considerable limitations, such as the complexity of fabrication, restricted cell migration and cell-cell interactions between different compartments, limited imaging accessibility, and a potential for causing cellular stress.

In this study, we introduce a simple approach for rapid prototyping of thermoplastic OOC devices by leveraging the hydrophilicity discrepancy among different regions within a microfluidic channel. Our strategy allows precise positioning of cells, ECM gels, and other fluid phases within the microchannel without the need for complex fabrication processes. As a proof of concept, we demonstrate that our OOC can support culturing of human hepatocytes for over 2 weeks and can be used for drug testing to assess the risk of DILI.

## Results

### Feasibility of using hydrophilicity discrepancy as the fluid guide

In contrast to the meniscus pinning effect provided by phase guide microposts (micropillars), an alternative approach to guide fluid flow is by leveraging the hydrophilicity discrepancy between different regions within the device. This method involves designing microfluidic channels with surfaces that possess contrasting hydrophilic properties: the more hydrophilic regions of the device attract and promote fluid flow, while the less hydrophilic regions repel and restrict fluid movement. To demonstrate the feasibility of this approach, we designed a two-channel device that contained one ECM lane for hydrogel patterning and cell embedding (either as single cells or as spheroids) and one medium lane for medium perfusion and vascularization during coculture. The chip fabrication involved several crucial steps, as illustrated in **Fig. 1A**. An ECM channel with a width of 300 μm and a medium channel with a width of 300 to 500 μm were designed with AutoCAD (**Fig. 1B**). Following this, pressure sensitive adhesive (PSA) tapes with a thickness of 120 μm were laser-cut to create the microchannel layer (PSA layer). Top and bottom substrates were prepared in tissue culture polystyrene (TC-PS), with holes drilled in the top substrate to facilitate engagement of fluid connectors. The PSA layer was aligned and adhered to the top substrate, and the tape covering the ECM lane was removed to allow surface functionalization via oxygen plasma treatment and polymer coating. The tape covering the medium lane was then removed. The PSA layer on the top substrate with a more hydrophilic ECM lane and a less hydrophilic medium lane was then bonded to the bottom substrate, after which the device was sterilized using ultraviolet (UV) or gamma irradiation. We selected polyethylene glycol (PEG) at 10 mg/ml with a molecular weight (MW) of 6 kilodaltons (kD) as the hydrophilic coating in our initial testing due to its relative inertness and its diverse utility in cell culture applications (*14*). When 1.5 μl of bovine collagen-I pre-gel at a concentration of 4 mg/ml was introduced to the ECM channel, a clear boundary between the ECM channel and the medium channel was observed, indicating successful establishment of surface hydrophilicity differences that supported hydrogel flow within the ECM channel (**Fig. 1C**).

**Figure 1.**
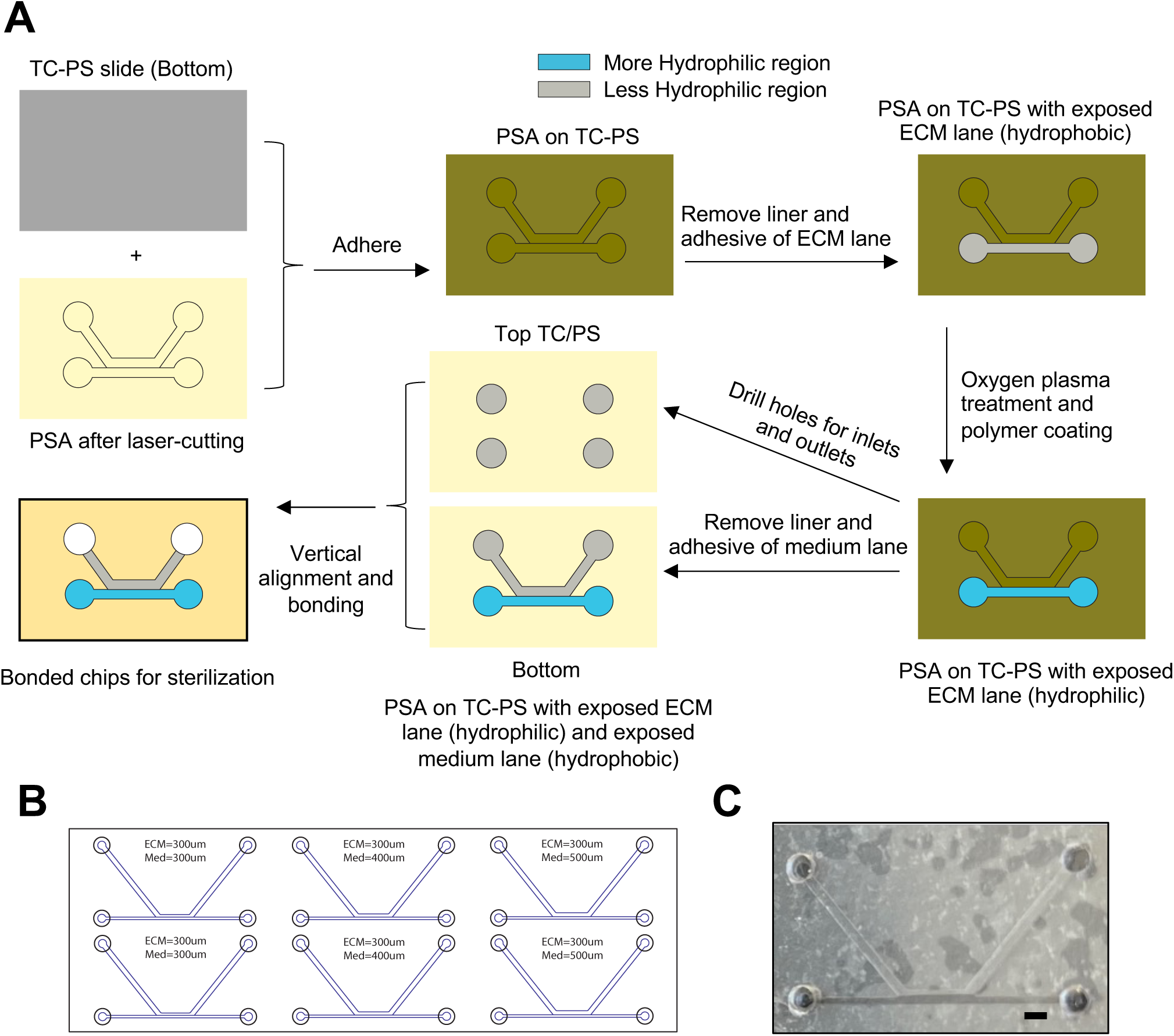
Feasibility of using hydrophilicity discrepancy as the fluid guide. **A)** Diagram showing the chip design and fabrication workflow. **B)** Layout and dimensions of the chip design. **C)** Photographs showing the containment of 4 mg/ml collagen-I gel within the ECM lane when only the top substrate was coated with PEG. Scale bar: 1 mm.

We then tested the effect of three additional coating configurations on hydrogel flow behavior. Flow testing of collagen-I pre-gel at 4 mg/ml showed that hydrogel solution could be confined within the ECM lane when the top or bottom substrate was selectively coated, but not when the entire top substrate was coated (**Fig. S1A, Fig. S1B #1-3**). In addition, we found that the confined flow of hydrogel solution within ECM lane was not affected by gamma sterilization (**Fig. S2**). We also evaluated different substrate materials, including polymethyl methacrylate (PMMA), PC, and glass, and found that they did not significantly change the flow behavior (**Fig. S3A, S3B**), nor when the channel height was increased to 240 μm and PEG coating was applied only to the bottom substrate (**Fig. S3C, Fig. S1B #4-5**). In summary, we demonstrated the feasibility of creating fluid guide by using PSA as a mask coupled with selective polymer coating to create regional hydrophilicity discrepancy.

### Optimizing polymer coating conditions

While successful hydrogel confinement within the ECM channel was achieved, the flow rate from one end of the microchannel to the other fell within the range of minutes. Such long durations could potentially lead to premature gelation before the ECM front reaches the other end, resulting in underfilling. This may also limit applications where higher concentrations of hydrogels are used. To address this issue, we sought to optimize the conditions for polymer coating. We conducted a comparative analysis of surface wettability of water on PS films at 2-, 24-, 120-, and 360-hour post-coating with PEG, polyvinylpyrrolidone (PVP), or polyvinyl alcohol (PVA) at various concentrations and MW. PVP coating at 1% or 2% significantly increased the water spreading area compared to PVA or PEG coatings, suggesting a higher degree of hydrophilicity with PVP coating (**Fig. 2A**). In addition, 1% PVP coating demonstrated stable surface functionalization within the PVP-coated group from 2 hours to 15 days post-coating under dry storage conditions, suggesting PVP as a more suitable choice than PEG or PVA for selective coating on the chip. To confirm this, we fabricated open-top devices (without top TC-PS substrate) using four different polymer coatings and compared their performance in guiding gel flow. In consistency with the film testing results, devices coated with 1% PVP displayed faster filling (less than 5 seconds) and fewer cases of chips exhibiting underfilling or overflowing along the ECM-medium interface (**Fig. 2B**).

**Figure 2.**
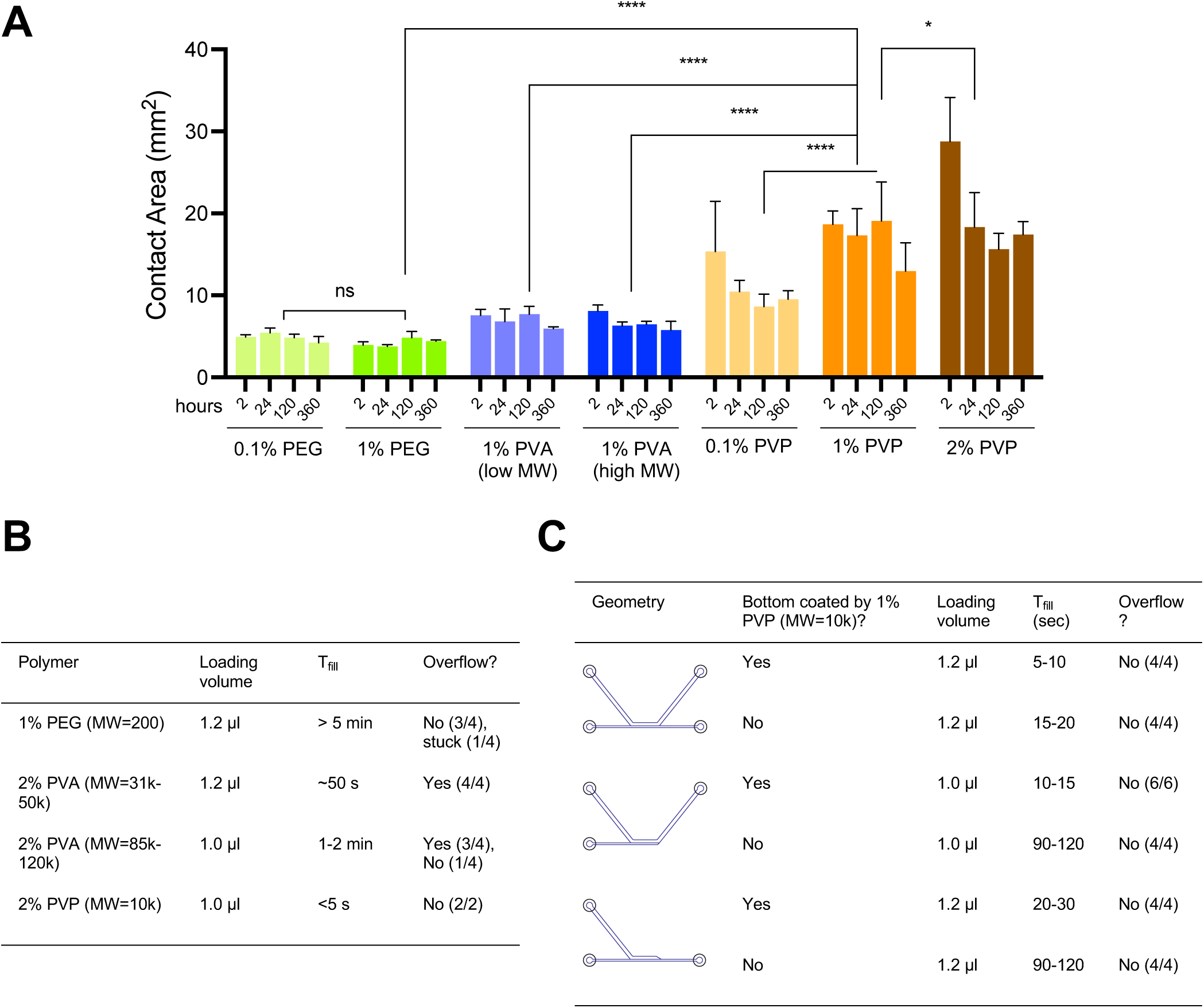
Optimizing polymer coating. **A)** Bar graphing comparing the contact areas of water dispensed on PS film at 2 hours to 360 hours post-coating with different types of polymers. **B)** Summary of the effect of different polymer coating on hydrogel flow behavior. Time required for the sample to complete fill the entire channel (t_fill_) is used to evaluate channel surface wettability and flow path is used to evaluate if selective coating on ECM channel can prevent the overflow of collagen-I solution into medium channel. **C)** Summary of the effect of channel geometry on hydrogel flow behavior.

We further validated these observations using open chips of three different geometries (**Fig. 2C**), all of which were coated with 1% PVP at the top PS substrate ECM lane. Despite differences in filling times, all chips demonstrated a normal flow behavior, without underfilling or overflow. Notably, when coating was applied to the bottom substrate, the time for gel filling was further reduced (**Fig. 2C**). Among the tested geometries, the “K”-shaped channel displayed the quickest gel filling (5-10 seconds) when both top and bottom substrates were selectively coated by using 2 layers of PSA (**Fig. 2C and Fig. S1B #6**). Thus, we have identified that the optimal polymer coating condition requires 1% PVP, which enables suitable functionalization of microchannel walls to create a precise hydrogel barrier.

### Modifications to increase chip performance and reliability for cell culture

Building upon the identified optimal coating conditions, we proceeded to test the feasibility of fabricating an integrated, perfusable OOC system suitable for cell culture. This system involved embedding cells within a hydrogel in the ECM lane, while using the alternate channel for medium perfusion (**Fig. 3A**). To facilitate fluidic connection between the chip channels and tubing, we used PMMA to construct a connection layer, which was bonded to the PS top substrate layer using a PSA (3M GPT-020F). The resulting five-layer device (PS-PSA-PS-PSA-PMMA) was mounted on a standard histology glass slide in a set of three to ease handling and imaging (**Fig. 3B, 3C**). Considering the multiple layers of the assembled chips, we explored the possibility of using UV radiation as a cost-effective alternative to gamma radiation for device sterilization. We subjected both top and bottom PS substrates, along with the exposed ECM lane, to UV dosages ranging from 0.015 J/cm^2^ to 1 J/cm^2^ within a UV-crosslinking device. Collagen-I pre-gel flow test revealed that UV exposure within this range did not significantly affect the flow behavior or gel filling time (**Fig. S4A, S4B**). To evaluate the sterilization efficiency as a function of UV intensity, we used a bacteria growth assay where 96-well plates containing serial dilutions of *Escherichia coli* (*E. coli*) were exposed to 0.3 J/cm^2^ or 1 J/cm^2^ of UV without cover or with the plate-lid or with the same 190-μm-thick PS film used as the chip top and bottom substrates. Analysis of bacterial colonies in inoculated agar plates found that only an exposure of 1 J/cm^2^ significantly decreased bacterial counts at concentrations ranging from 10^5^ to 10^9^ plaque forming unit (pfu)/100 μl (**Fig. S4C**). Therefore, we concluded that complete sterilization requires direct UV exposure of all surfaces surrounding the cell culture chamber at an energy of at least 1 J/cm^2^.

**Figure 3.**
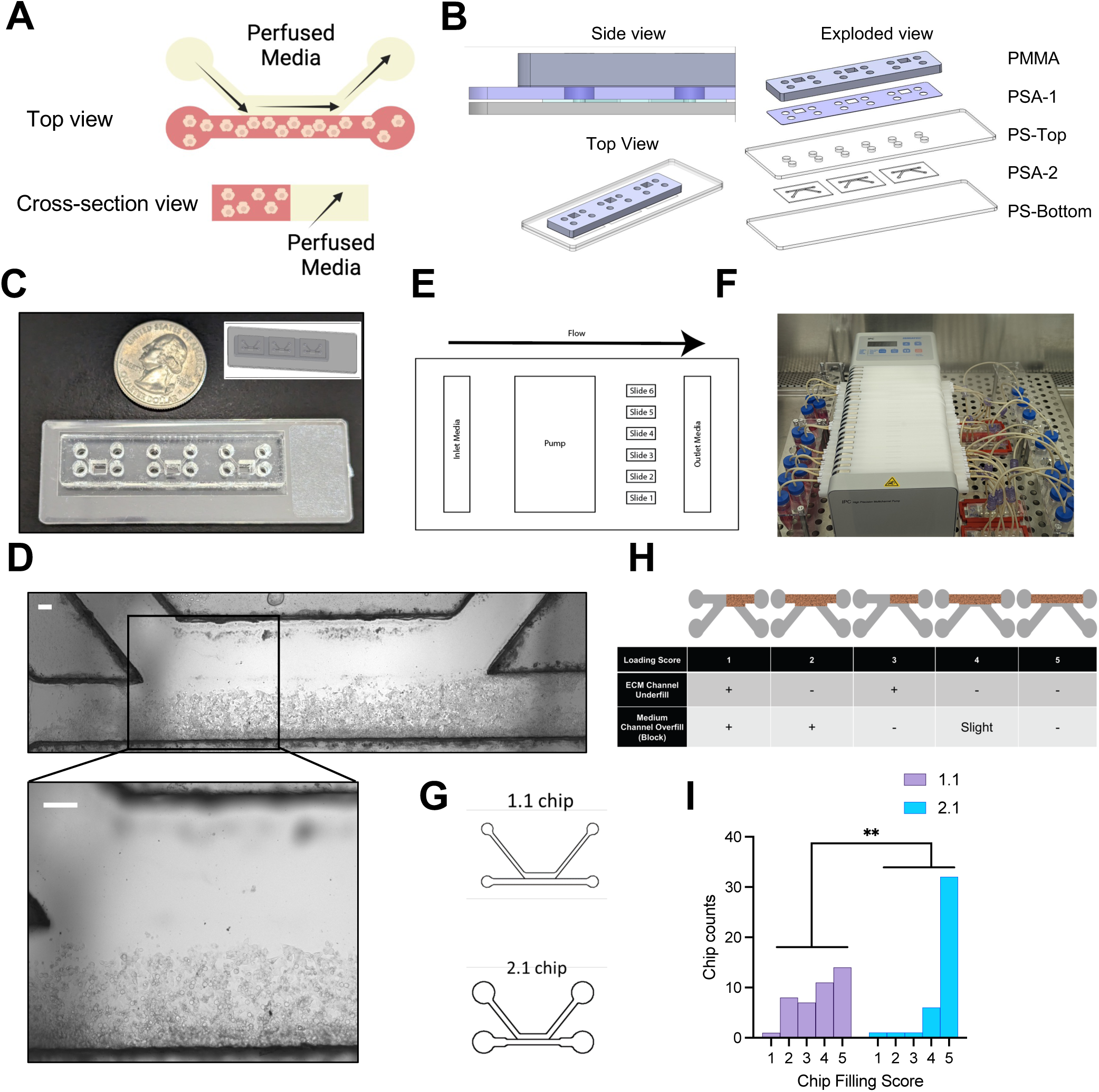
Integrated Chips with perfusion for cell culture. **A)** Cartoon showing the embedding of cells in the ECM channel and the perfusion of culture medium in the medium channel. **B**) Schematic views of the 5-layer device from top to bottom: PMMA, PSA-1, PS top substrate, PSA-2, PS bottom substrate. **C**) Photographs showing a set of 3 assembled chips on a glass slide. **D**) Phase contrast images showing distribution of HepG2 cells within the ECM channel. Scale bar: 50 μm. **E**) Diagram showing the arrangement of different components in the perfused chip culture system. **F**) Photograph showing the setup of an experiment on a standard tissue culture incubator. From left to right: inlet reservoirs, peristatic pump, chips on slides sitting on a chip carrier, and outlet reservoirs. **G**) Diagram showing the original chip design (1.1 chip) and the design with enlarged ECM channel ends. **H**) Diagram showing the chip filling score. A score of 4 or 5 is considered acceptable for downstream culture. **I**) Histogram of chip filling scores for 1.1 chip and 2.1 chip. N= 41 for each group. Fisher’s exact test, **p<0.01.

We subsequently assessed the suitability of our device for seeding cells embedded in the hydrogel. HepG2 liver cells, with a density of 5 x 10^7^ cells/ml, were suspended in a 2.56-mg/ml solution of human collagen-I pre-gel and incubated at 37 °C for 1 hour to facilitate gelation. Consistent with results from experiments using only gel, we found that the gel-cell mixture remained precisely confined within the ECM channel of the device (**Fig. 3D**). We then used a 24-channel peristatic pump to drive unidirectional medium flow in the medium channel (**Fig. 3E, 3F**). Even at a flow rate of 100 μl/hour, corresponding to a shear stress of 0.033 dyne/cm^2^, for at least 5 days that we tested, the solidified collagen gel and embedded cells remained intact (**Fig. S5**). For subsequent experimentation, we selected a flow rate of 30 μl/hour for ease of sampling.

Despite the impressive gel flow behavior within the channel, we encountered challenges with the fabrication process, specifically incomplete peeling-off of the PSA liner, leading to partial channel blockage. To address this, we designed a new chip (2.1 chip) with a medium channel width of 500 μm as opposed to the original 300 μm (1.1 chip) and expanded the width of both ends of the ECM channel from 500 μm to 800 μm (**Fig. 3G**). This modification made peeling-off the PSA liner much easier, significantly reducing the percentage of chips with debris that could potentially block fluid flow, from 29% to 3% (**Fig. S6**). To better evaluate the flow response of cell-laden hydrogels, we developed a semi-quantitative scoring system (**Fig. 3H**). An analysis of over 40 chips revealed that the modified chip design significantly increased the percentage of chips that met the criteria for downstream perfusion and cell culture. Specifically, the percentage of chips with a filling score of 4 or 5 rose to 92.7%, a significant improvement from the 61.0% achieved with the original design (**Fig. 3I**).

### Liver Chip model characterizations

Based on these modifications that boosted the performance and reliability of the chips, we investigated the potential of our device for sustaining long-term cell culture with selective coating on both top and bottom substrate ECM lanes (configuration shown in **Fig. S1B** #5). HepG2 cells were cultured at a density of 5E+07/ml in a 2.56-mg/ml human collagen-I gel for a period of 14 days, under a constant flow rate of 30 μl/hour. Immunofluorescence staining showed a high level of albumin expression in these cells (**Fig. 4A**). Quantifying albumin levels in the medium outflows by enzyme-linked immunosorbent assay (ELISA) also revealed increased albumin production on-chip, reaching up to 15 μg/1E+06 cells/day, which is significantly higher than those observed in two-dimensional (2D) culture (**Fig. 4B**). In summary, our device is capable of supporting long-term cell culture in a three-dimensional (3D) environment under dynamic perfusion, with hepatocytes displaying enhanced metabolic competence compared to their 2D counterparts.

**Figure 4.**
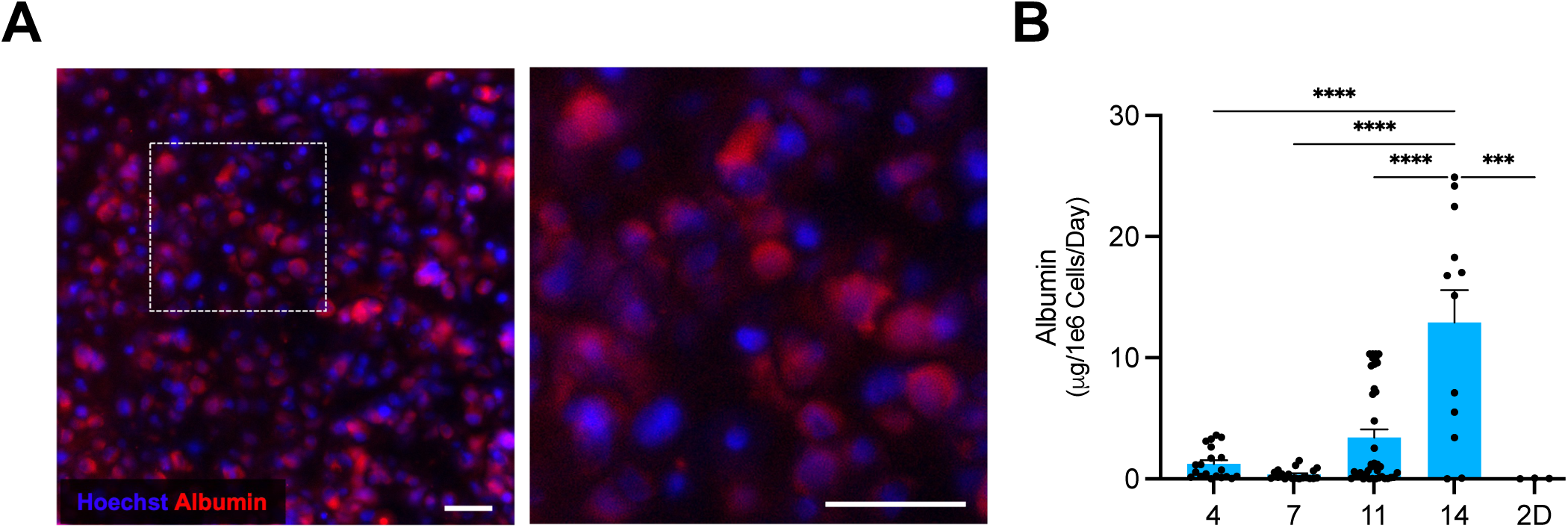
Characterizations of the Liver Chip Model. **A)** Immunofluorescence staining of albumin of the cultured HepG2 cells in the liver chip at day 11 of culture. Scale bar: 50 μm. **B**) Albumin secretion at day 4, 7, 11, and 14 of culture on chip compared with 2D control. Data in **B** represent mean ± SEM; N=2, n = 3-21 biological replicates (individual chips) per group; one-way ANOVA with Tukey’s post-hoc multiple comparisons correction, ***p<0.001, ****p<0.0001.

### Feasibility of liver chip model for DILI testing

One advantage of OOC made from thermoplastic materials compared to those made with PDMS is their non-absorbency of hydrophobic molecules, thus making them a superior choice for applications such as assessing drug-associated liver toxicity. To determine whether our device is suitable for DILI testing, we utilized troglitazone, a diabetic drug withdrawn from the market due to severe hepatotoxic effects in humans that could not be predicted from regulatory animal or *in vitro* studies (*15*). We introduced troglitazone into the medium channel at seven-fold (7x) or twenty-fold (20x) of its maximum plasma concentration (C_max_) on day 11 of the culture (**Fig. 5A**). Medium was collected 72 hours later, and chips underwent CellPainting, a high-content image-based assay for morphological profiling using multiplexed fluorescent dyes (**Supplementary Table 1**) (*16*). Analysis of secreted albumin from the medium outflows indicated that troglitazone at 20x C_max_ decreased albumin production, while a minor effect was observed at 7x C_max_ concentration (**Fig. 5B**). CellPainting images also showed that high concentration of troglitazone led to a reduction in cell count and correlated with distinct morphological changes in the structure and intensity of mitochondria and the cytoskeleton (**Fig. 5C**). We then extracted image features at the cellular level and performed dimension reduction to visualize and compare morphological alterations across treatment groups. Uniform Manifold Approximation and Projection (UMAP) shows more drastic changes in cell morphology at 20x C_max_ concentration than 7x C_max_ concentration, consistent with the albumin assay result. We further ran a random forest (RF) classifier to explore how well the different treatment groups were separated and which features might be driving the separation. We split the data into a training set and a test set. The RF classifier reached a ROC (Receiver Operator Characteristic) AUC (area under the curve) score of 0.93 for the test set. Feature importance analysis revealed that top-ranking features are intensity features, indicating an impact of troglitazone on the binding of CellPainting dyes to organelles (**Fig. S7**). These results suggest that our liver chip device could be integrated with biochemical assays or high-content imaging assays for DILI risk prediction.

**Figure 5.**
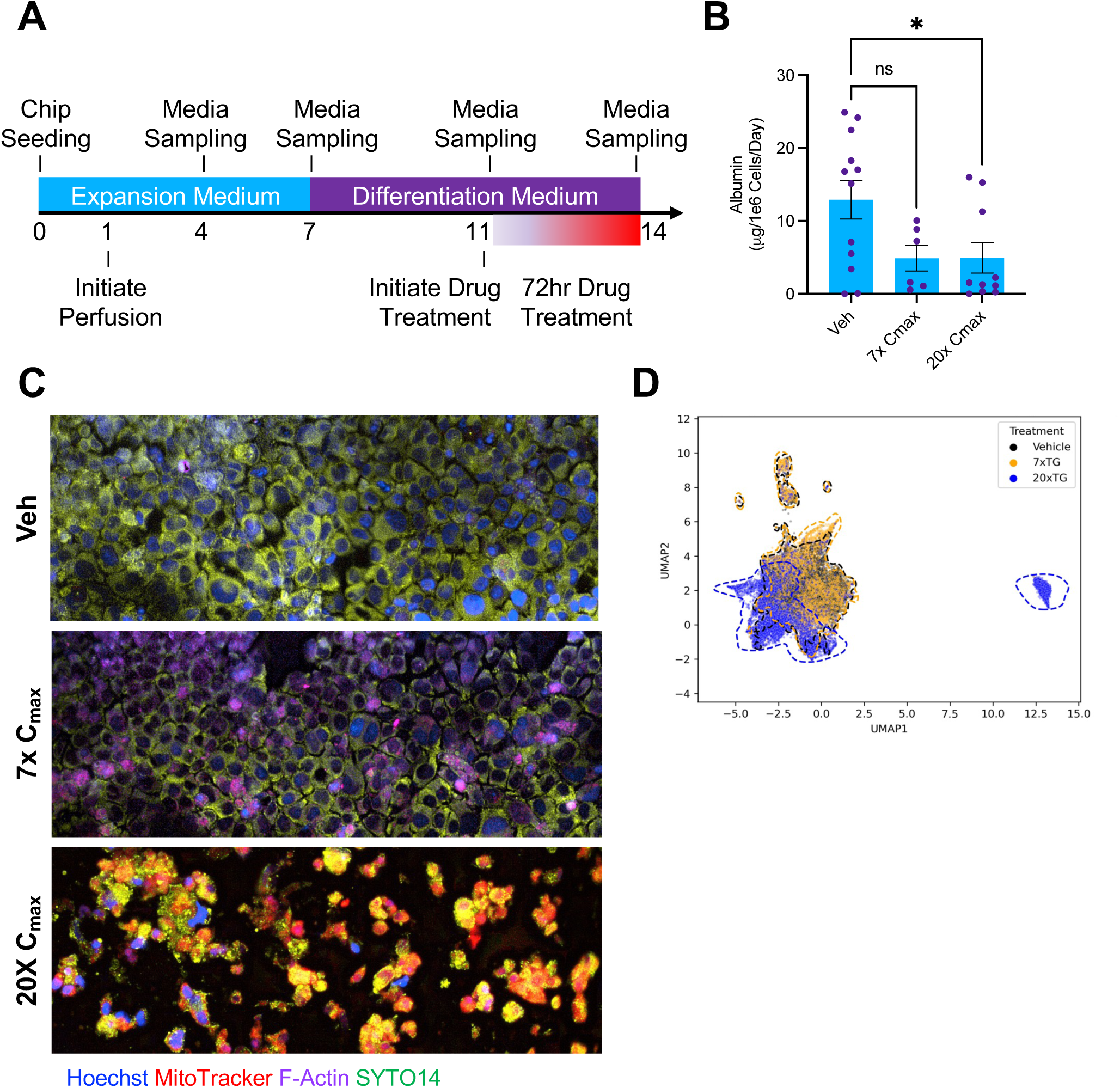
Troglitazone toxicity and morphological analyses. **A)** Schematic diagram showing the workflow of cell culture and drug treatment on the liver chip for DILI assessment. **B**) On day 11, On-Chip cultures were treated with either Troglitazone (TG, 7x C_max_ or 20x C_max_) or Vehicle (0.2% DMSO) and albumin was measured from the outflow at 72 hours post-treatment. Data in **B** represents mean ± SD; N=2, n = 3-8 biological chip replicates; one-way ANOVA with Tukey’s post-hoc multiple comparisons correction, ns, not significant. **C**) Following 72 hours of drug treatment, cells on-chip were subjected to the CellPainting assay. Representative data shown, N=3, n=2-3. Stained devices were imaged at 20x. Scale bar: 50 μm. **D**) UMAP of cellular features showing phenotypic clustering with troglitazone treatment, noting distinct differences in object-level metrics among the treatment groups. N=2, n=1-2 per group.

## Discussion

In this study, we presented a unique approach to fabricating OOC devices using thermoplastic materials. By generating a hydrophilicity discrepancy within the device, we were able to guide fluid flow, thus creating a Liver Chip Model that enabled long-term 3D cell culture and assessment of DILI risk.

The use of soft lithography with PDMS has opened up a multitude of applications in biomedical engineering (*17, 18*), including the OOC field. While PDMS remains the preferred material due to its biocompatibility, optical transparency, and gas permeability (*8*), it also carries significant drawbacks such as high mass-production costs and drug adsorption/absorption. To overcome these limitations, recent studies have explored the use of thermoplastics due to their optical transparency, cost-effectiveness, low drug adsorption/absorption, and scalability for mass production (*8, 19–22*). However, prototyping complex designs with thermoplastics has been challenging.

Our strategy differs from conventional approaches to create multi-channel OOC devices. By combining PSA and selective polymer coating, we were able to create a contrast in surface hydrophilicity behaviors, which resulted in distinct cellular compartments within the OOC device. Notably, this technique does not require the introduction of physical barriers, such as micropillars or microposts (*2, 8, 12, 23*) or intricate handling procedures (*24*), thereby simplifying the fabrication process and reducing costs. Our study demonstrates that OOC designs of various thermoplastic materials, channel designs, and microstructures can be prototyped facilely, which is amenable to rapid prototyping. Our approach differs from other membrane-free designs of OOC that mainly relies on the meniscus pinning effect offered by phase guide microposts or micropillars (*2, 8, 12, 23*), which still have a partial intermittent barrier, typically made of PDMS on top of glass, along the compartment interface.

Our Liver Chip Model exhibited enhanced hepatocyte functions in comparison with standard two-dimensional culture. This is consistent with previous publications suggesting that dynamic 3D culture improves hepatocyte long-term viability and metabolic competence (*25–27*). Our model also demonstrated the cytotoxic effects of troglitazone, a known hepatotoxic drug, suggesting the potential of our platform for pharmaceutical applications such as large-scale DILI testing, especially when combing it with high-content imaging assays, such as CellPainting, and morphological analysis powered by machine-learning algorithms.

We identified PVP as a suitable polymer for selective coating. PVP is a water-soluble, biocompatible, and non-toxic polymer that has been used as a supplement in media, as a coating on various surfaces, and as a reinforcing material in tissue engineering (*28*). While the risk of PVP on cell viability is low, whether it affects long-term cell function requires further investigation. In addition, future work is also warranted to identify optimal coating conditions that balance pre-gel filling time and the risk of gel overflow for hydrogels with different concentrations or viscosities.

Our study has certain limitations. For instance, the use of PSA during the fabrication process might induce cytotoxic effects that influence long-term cell viability. Additionally, potential autofluorescence might interfere with imaging assays. However, these issues could potentially be resolved by using different types of PSA or replacing it with other thermoplastic materials during mass production. Additionally, our study centered on the engineering strategy and primarily used the HepG2 cell line. Despite successful long-term cell culture and enhanced metabolic function compared to 2D culture, it is crucial to validate our system using primary hepatocytes and integrating liver sinusoidal endothelial cells along with other nonparenchymal cells. This will increase the model complexity, better mimicking the *in vivo* multicellular environment and potentially improving prediction accuracy in drug testing applications, such as DILI.

In summary, our research introduced a novel approach in the design and application of OOC models by leveraging hydrophilicity discrepancy within the device. Our study confirmed that this unique strategy is not only feasible but highly effective in enabling precise hydrogel patterning and compartmentation of cell culture environment. This approach provides new opportunities for the development of high-throughout and reduced-cost OOC models with potential applications in drug screening and personalized medicine.

## Materials and Methods

### Chip design and fabrication

The chip fabrication process involved three key steps: construction of the top substrate with a PMMA cap, polymer coating on the ECM channel of the top substrate, the bottom substrate, or both substrates as specificized in each graph, and chip assembly and sterilization. First, the PMMA cap was fabricated using a Glowforge Pro laser cutter, with specific parameters set for the cutting process: medium clear acrylic material, a speed of 170, full power, a proof grade cut mode, and one repeat. The top substrate was drilled, cleaned, and then affixed with the PMMA cap using a Darwin Microfluidics fluid connector and a 3M PSA. The top substrate was cleaned once more, ensuring the removal of any residual materials. PSA pieces were then attached to the frosted front side of the substrate, aligned with the drilled holes. The substrate then underwent a corona treatment using a plasma cleaner (Harrick Expanded Plasma Cleaner, 115V, #PDC-001). Post-treatment, the substrates were immersed in polymer solution, blown dry, and heated on a hotplate (VWR, #76447-044) to ensure complete drying. The following polymers were tested in this study: 1% (w/v) PVP (MW=10,000, MilliporeSigma, #PVP10-100G), PEG (MW=6,000, MilliporeSigma, #8074911000), PEG (MW=200, MilliporeSigma, # U214-07), PVA (87.0-89.0% hydrolyzed, average MW 31,000-50,000, Thermo Fisher, #183141000), and PVA (88% hydrolyzed, average MW 85,000 to 120,000, Thermo Fisher, #450140250). Subsequently, the adhesive layer and plastic backing were removed to expose the medium lane of all channels. The top substrate and bottom substrate underwent UV sterilization. The substrates were placed into a Stratalinker UV-crosslinker, which was set to an energy level of approximately 1 J/cm^2^. Post-irradiation, the substrates were covered and transferred into the biological safety cabinet (BSC) for chip assembly. The device was assembled by adhering the PSA layer of the capped top substrate onto the front surface of the bottom substrate, ensuring that as many air pockets in the PSA layer are removed as possible, particularly near the channel walls. The assembled device was then sterilized again with UV irradiation at the same energy level as before. The process was repeated for all devices, with sterilization measures in place at all times to prevent contamination.

### Measurement of liquid contact area

A droplet test was performed to evaluate contact angle and surface hydrophilicity. A 2 µL volume of distilled water was dispensed on the PS film at different time points post polymer coating. Images were taken and surface contact area was measured using Fiji (ImageJ) for each droplet. At least 3 droplets were measured and averaged for each film.

### Hydrogel flow test

To prepare fresh 4 mg/mL of collagen I hydrogel precursor solution, 8 µL of deionized water, 10 µL of 10x PBS, 2 µL of 1-M NaOH, and 80 µL of 5-mg/ml stock bovine collagen I solution were mixed and stored on ice. Alternatively, stock solution of 0.1-M NaOH was prepared in 5x PBS and mixed with 5 mg/mL of bovine collagen at a 1:4 ratio. For hydrogel flow testing on the open chips, device was placed on an ice bath and loaded carefully with 5 µL of hydrogel solution. The flow behavior was observed and the flow time from gel entry to completion was recorded. Images were captured quickly when the flow was complete. Subsequently, the device was placed in a standard tissue culture incubator at 37 °C to allow gelation for 30 minutes and images were taken again. Similarly, for hydrogel flow testing on the assembled chips, 4 mg/mL of collagen I solution was prepared fresh and loaded to the inlet port of the ECM channel at a volume from 1.25 µl to 2 µl.

### Cell culture

HepG2 (CRL-3216) cell line were obtained from American Type Culture Collection (ATCC). HepG2 cells were cultured in DMEM (Gibco, Cat# c11995500BT) supplemented with 10% (v/v) fetal bovine serum (PAN, Cat# ST30-3302), 100 U/mL of penicillin, and 100 µg/mL of streptomycin at 37 ℃ in 5% CO_2_. Cells were regularly checked for free of mycoplasma contamination.

### 3D cell culture on chip

On day 1 of cell culture on chip, Sterile Five-O™ 5ml MacroTubes (Fisher Scientific, #501929153) were used as inlet and outlet medium reservoirs. Holes were punched in the lid using a hammer-driven hole punch (McMaster-Carr, #3424A13). The following tubing, connectors, and plugs were organized into autoclave bags and autoclaved: male Luer Lock to 1.6-mm Barb Connector (2x per chip, Qosina, #11735), stainless steel dispensing needle with Luer Lock connection. ½” needle length, 27 gauge (1x per chip, McMaster-Carr, #75165A688), male Mini Luer Fluid Connectors (2x per chip, Darwin Microfluidics, #CS-10000095), male Mini Luer plugs (2x per chip, Darwin Microfluidics, #CS-10000054), Masterflex Ismatec Pump Tubing, PharMed BPT, 1.02-mm ID; 100 ft (3x per chip [15 mm and 25 mm], VWR, # MFLX95809-28), Masterflex Ismatec Pump Tubing 0.25 mm (or 0.19 mm, 1x per chip PharMed BPT, #MFLX95723-12). On day 0, all components were arranged in the BSC and connected in the following order: for inlet line, connecting shorter tubing segment to Luer, to needle, to two-stop tubing, to needle, to Luer, to shorter tubing segment, and to Darwin Mini Luer connector; for outlet line, connecting Darwin connector to longer tubing to the outlet reservoir. Male Mini Luer Plugs (2x per chip, Darwin Microfluidics, #CS-10000030) were inserted into the ECM channel on both ends post gelation to prevent evaporation. Human collagen-I (Advanced Biomatrix, #5007) was prepared according to manufacturer’s instruction. HepG2 cells were harvested and resuspended in chilled collagen solution at a final density of 5E+07 cells/ml. 1.35 µL of cell suspension was pipetted into the ECM channel inlet port. Chips were placed in square petri dish containing 2 ml of water in a 50-ml tube lid to prevent evaporation during gelation. Chips were put away in the incubator (37 °C, 5% CO_2_) and incubated for 90 minutes to allow complete gelation. An equal volume of cell suspension was added to a 96-well plate as a control. Darwin plugs were inserted into the ECM channel following gelation to avoid channel evaporation. Chips were connected to the pump with 6 mL of cell growth medium added to the inlet reservoir tube. After priming to remove the bubble, flow was initiated using a multichannel peristaltic pump (VWR, #MFLX78001-40) with a flow rate of 30 μL/hour to 60 μL/hour (depending on tubing internal diameter). Outlet medium was collected every 72 hours and stored at −80 °C. Inlet medium was refilled to 6 ml during outlet medium sampling.

### Albumin and urea assays

Albumin assay was performed using the human albumin ELISA kit from RayBiotech (#ELH-Albumin-1) according to the manufacturers’ instructions. Briefly, frozen (−80 °C) effluent samples were thawed overnight at 4 °C prior to assay. Chip samples were used without additional dilution; static control samples were diluted 1:10 in the assay buffer. Absorbance at 450 nm was measured for albumin assay and optic density at 520 nm was measured for urea assay, respectively, both using the BioTek Synergy Neo2 microplate reader. All downstream assay data were normalized to µg/1E+06 cells/day format. Normalization was performed by assessing the cell count and perfused medium volume at each timepoint. Briefly, downstream assay output values at each timepoint were calculated as below, with X representing the downstream assay output, *C_Count_* representing the cell count at the indicated timepoint, *V_perf_* representing the perfusion volume in ml at the indicated timepoint, and *D_perf_* representing the total days of perfusion since the previous timepoint:

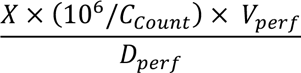

### Drug treatment

The drug dosing concentrations were based on the unbound human C_max_ of troglitazone (1.6 μM). For troglitazone, 20x C_max_ (32 μM) and 7x C_max_ (11.2 μM) were prepared as 500x stocks (366 mM and 5.6 mM, respectively) in sterile DMSO, aliquoted, and stored at −80 °C. 500x stocks of sterile DMSO served as a vehicle control, with a final 1x DMSO concentration of 0.2% for all treated and control samples. On day 11 of culture, inlet medium was refreshed with maintenance medium containing the indicated drug treatment C_max_ or vehicle control at a final concentration of 1x. On-chip cultures were perfused with drug- or vehicle-containing medium for 72 hours, prior to endpoint sampling and assessment.

### CellPainting assay

The CellPainting protocol (*16*) was modified to accommodate the on-chip 3D culture and performed *in situ* 72 hours post-drug treatment. Cells were rinsed with pre-warmed 1x PBS (Gibco, #10010023), and then incubated in Live-Cell staining solution (**Supplementary Table 2**) in the dark at 37 °C in 5% CO_2_ for 45 minutes. Following incubation, cells were rinsed with pre-warmed 1x PBS three times (with 10-minute incubation for each wash step throughout protocol) and fixed with 3.7% PFA (Thermo Fisher Scientific, #J61984.AP) for 20 minutes at room temperature. Fixed cells were washed twice with 1x PBS then permeabilized in 3D Permeabilization Buffer (0.5% Triton-X100 in 1x PBS) for 20 minutes. Following permeabilization, chips were washed twice with 1x PBS prior to adding the staining solution (**Supplementary Table 1**). Chips were incubated for 1 hour at room temperature, then washed three times with 1x PBS-T (1x PBS with 0.05% Tween20 and 0.02% sodium azide) and once with 1x PBS. Prior to imaging, antifade Imaging Buffer (0.1-mM VectaCell Trolox CB-1000-2 in 1x PBS) was added to chips to prevent photobleaching. Stained chips were stored at 4 °C in the dark for later use.

### Automated image acquisition and processing

Image acquisition was performed with the Agilent BioTek Lionheart LX Automated Fluorescent Microscope in 6 fluorescent channels provided in **Supplementary Table 2**. Gen5 software (Agilent/BioTek) was utilized to fully automate image acquisition, processing, and on-instrument cellular analysis. Each liver-chip was acquired by a 20x objective (Olympus Plan Fluorite 0.45NA #1220517) at 5 overlapping region of interest and 50 Z-slices to capture the entire ECM channel. Following acquisition, image processing steps included stitching, deconvolution, background subtraction, and Z projection using focus stacking.

### Morphological analysis

Cellular segmentation and high-content analysis was performed as part of the fully automated imaging protocol described above with the Gen5 software (Agilent BioTek). Individual cells were assessed for a number of intensity- and geometry-based metrics. Extracted object-level features were normalized with respect to the vehicle control chips and further batch-corrected using the sphering transformation (*29*). UMAP (*30*) dimension reduction was performed to visualize the 2D projections of Cell Painting features, showing clustering patterns in different treatment groups. We further split the data into a training set (70%) and a test set (30%). A Random Forest classifier was used to classify single cell features into different groups, with different treatments as classification labels. This classifier was trained on the training set and subsequently validated on the test set. The performance of the classification was primarily evaluated using the Area Under the Curve of the Receiver Operating Characteristics (AUC ROC) score. However, additional performance metrics such as accuracy, precision, recall, and F1 score were also assessed. Through this process, the RF classifier provided a feature importance ranking, thereby elucidating the features that predominantly contribute to the differentiation of classes.

### Statistical analysis

All experiments were repeated at least two times. Data are displayed as mean values ± standard deviations (SD) unless otherwise noted. Graphing and statistical comparison of the data were performed using GraphPad Prism 9.0. Two-group comparisons were assessed using the two-tailed Student’s t test; comparison of three or more groups were analyzed by one-way ANOVA with Tukey’s multiple comparisons test. *p* values < 0.05 were considered to be statistically significant; * p< 0.05; **p< 0.01; ***< 0.001; n.s., not significant. Due to the sample variation for on-chip culture, a ROUT analysis was performed to remove sample outliers for Fig. 4B and Fig. 5B with a Q value of 5 (corresponding to a maximum false discovery rate of 0.05).

## Author Contributions

X.X., H.B., X.Q., Y.S.Z., and M.P. conceptualized the study. H.B. Drafted the manuscript with help from all authors. X.X., M.P. and H.S. designed the chip and other accessories. M.P. and T.M. fabricated the device and other accessories. M.P. performed gel flow testing. K.N.O., H.B. performed cell culture on chip. P.G. and YC.Y. conducted albumin, urea, and CellPainting assays. YC.Y. and J.M., K.N.O. conducted imaging analysis. L.S., D.H., S.M., C.L., and Y.S.Z. provided feedback during the study and during manuscript revision. H.B., and K.N.O. performed data analysis. H.B. and X.X. finalized the manuscript. The final version of manuscript has been approved by all authors.

