## Supplementary Information for "Rapid Prototyping of Thermoplastic Microfluidic 3D Cell Culture Devices by Creating Regional Hydrophilicity Discrepancy"

### Supplementary Figure 1

**A**

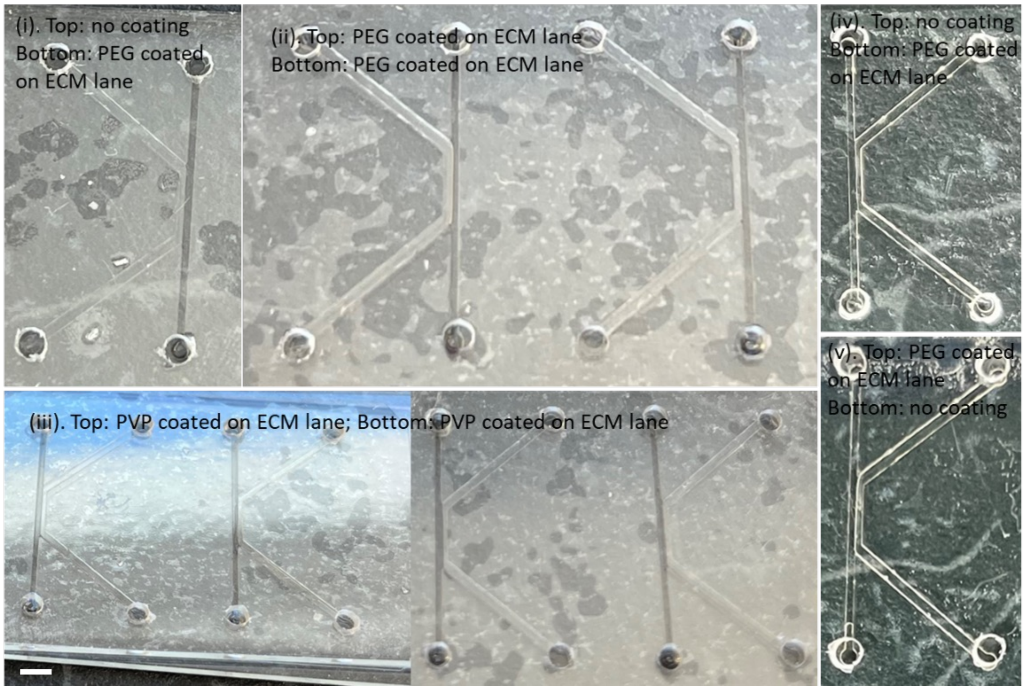


**B**

| # | Substrate Combination | Medium Channel | ECM Channel | Merged Channel | Overflow? |
|---|---|---|---|---|---|
| 1 | Top / Bottom | | | | No |
| 2 | | | | | No |
| 3 | | | | | Yes |
| 4 | | | | | No |
| 5 | | | | | No |
| 6 | | | | | No |

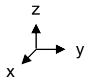


More Hydrophilic region
Less Hydrophilic region

**Supplementary Figure 1. Effect of different coating configurations on hydrogel flow behavior. A**) Photographs shows flow test results (before gelation) using chips (channel height: 120 μm) with different coating configurations using (i-iii). ~1.5 μL 0.35% rat collagen I solution; and (iv-v). ~1.5 μL 0.4% bovine collagen I solution, as hydrogel solution. Scale bar: 1 mm. **B**) Summary of hydrogel flow test using chips with different coating configurations.

### Supplementary Figure 2

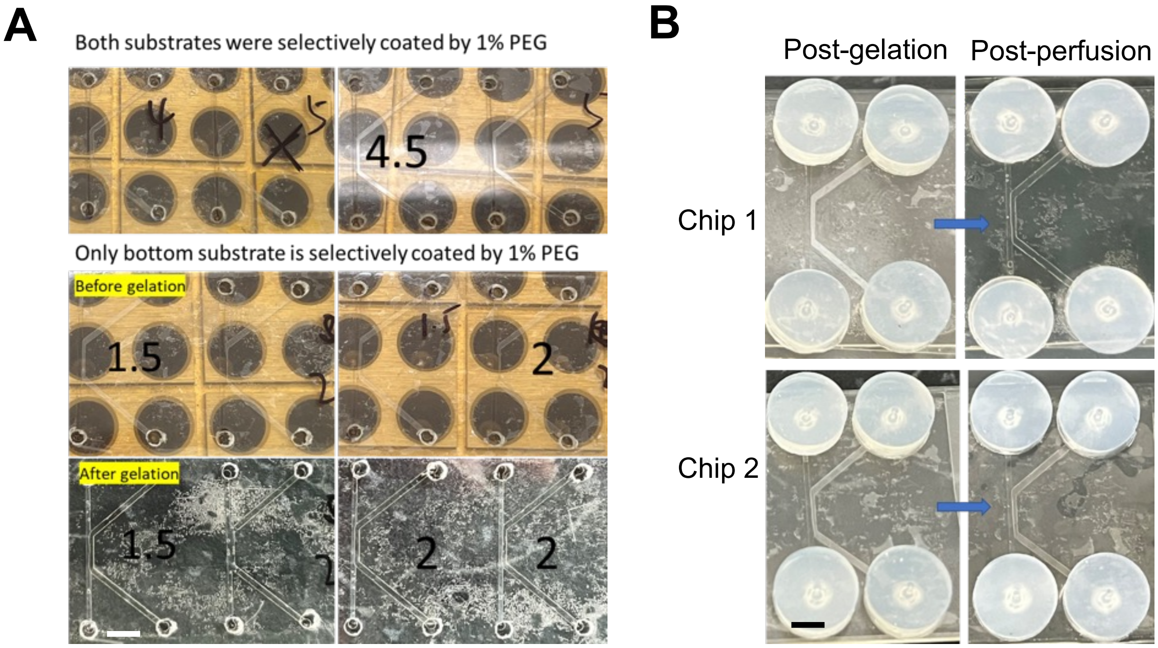


**Supplementary Figure 2. Effect of gamma irradiation on hydrogel flow behavior. A)** Photographs showing results after loading 0.4% bovine collagen I solution into chips after sterilization by gamma irradiation at 25 kilogray (kGy). Numbers in the images are loading volumes. Scale bar: 2 mm. **B**) Photographs showing flowing and gelation results after loading ~1.5 µL 0.4% bovine collagen I solution into chips after sterilization by gamma irradiation at 25 kGy (channel height: 120 µm). The tubing connectors were adhered to the inlets and outlets between gel loading and gelation steps. After gelation the resulting chips are ready for perfusion experiments by flowing culture medium solutions across medium lane using a peristaltic pump. Scale bar: 2 mm.

### Supplementary Figure 3

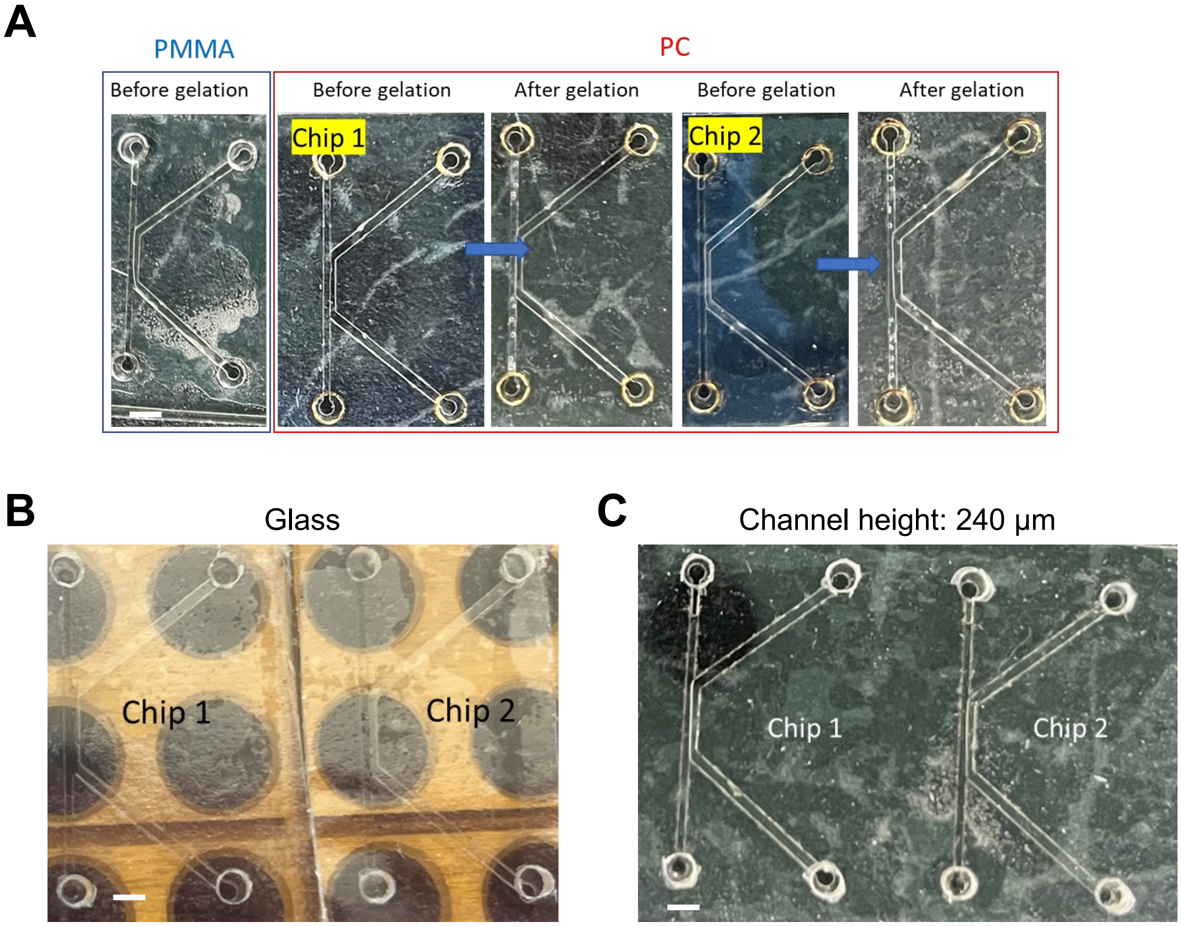


**Supplementary Figure 3. A&B) Effect of substrate materials and channel heights on hydrogel flow behavior.** Photographs showing flowing and gelation results after loading 0.4% bovine collagen I solution into non-sterilized chips (channel height: 120 µm) with different types of plastic substrates. **A)** The top substrate was either PMMA or PC slide with ECM lanes coated by 1% PEG, the bottom substrate was pristine TC-PS slide without surface modifications. **B**) The top substrate was TC-PS slide with ECM lanes coated by 1% PEG, the bottom substrate was glass slide with ECM lanes coated by 1% PEG. **C**) The top substrate was TC-PS slide with ECM lanes coated by 1% PEG, the bottom substrate was TC-PS slide with ECM lanes coated by 1% PEG (channel height: 240 µm). Scale bar: 2 mm.

### Supplementary Figure 4

## A

UV dosage: 0 | 0.015 J/cm$^2$ | 0.3 J/cm$^2$ | 1 J/cm$^2$

Gelation

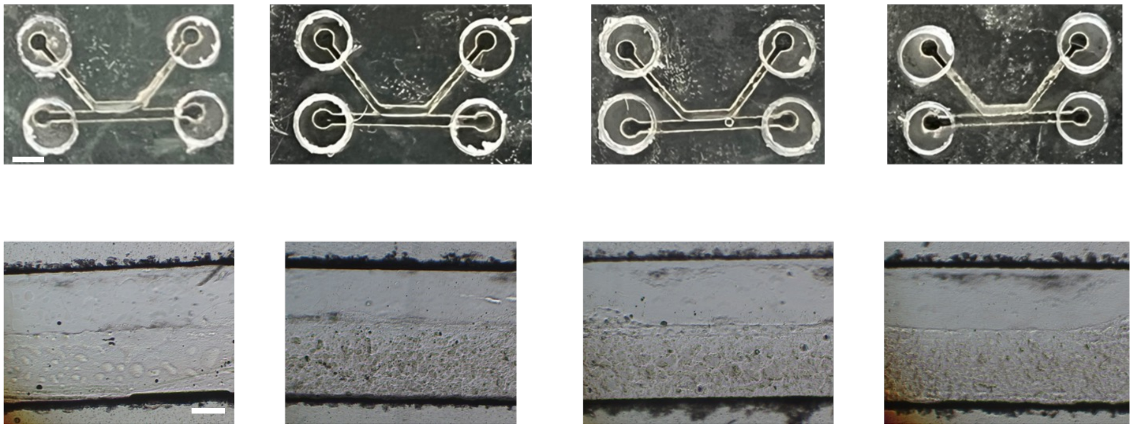

## B

| Geometry | UV dosage | Load volume | $t_{fill}$ | Overflow? |
|---|---|---|---|---|
| 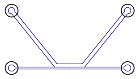 | 0 | 1.2 uL | 5-10s | No. 4/4 |
| Bottom substrate coated? Yes | 0.015 J/cm$^2$ | 1.2 uL | 10-20s | No. 2/4<br>Overload 2/4 |
| | 0.3 J/cm$^2$ | 1.2 uL | 5-10s | No. 4/4 |
| | 1 J/cm$^2$ | 1.2 uL | 5-10s | No. 4/4 |

## C

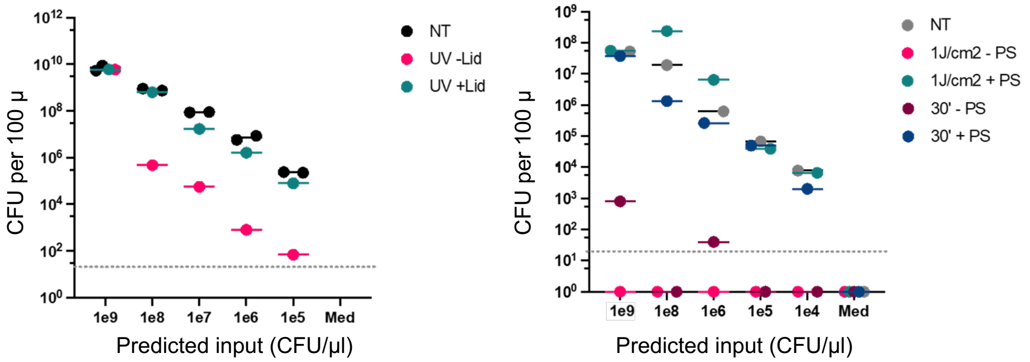

**Supplementary Figure 4. Effect of UV exposure on gel flow behavior and sterilization efficacy. A)** Photographs showing the results of hydrogel flow tests where the ECM channel of the top substrate and the entire bottom substrate were coated by PVP. Both substrates were sterilized by UV at 254 nm with different energy levels. In all cases, the hydrogel solution only filled the ECM channel without significant overflow into the medium channel. Scale bars: 2 mm (top) and 250 μm (bottom). **B)** Summary on the effect of UV sterilization at different energy levels on hydrogel solution flow behavior in Mercury chips. The entire bottom substrate was pristine without coating before UV sterilization. Note for results shown in the table, when hydrogel solution was overflowed, it filled the entire ECM and medium channel. **C**) Plots showing measured cell concentration from plate-well treated under various sterilization conditions, including different times or energy levels of UV exposure, whether the plate-well was covered PS film was covered during UV exposure.

#### Supplementary Figure 5

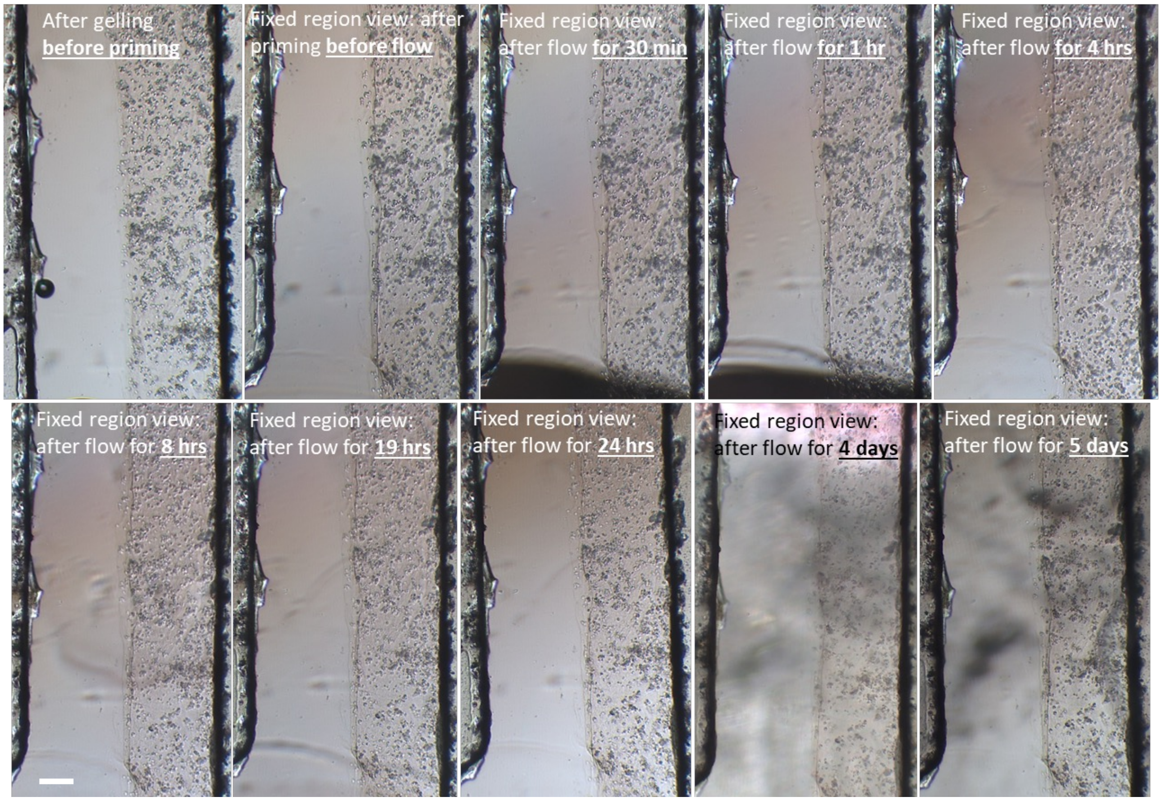


**Supplementary Figure 5. Gel Stability under perfusion**. Optical microscope images showing gel and cell morphology under continuous flow of culture medium at 100 µl/hour. The chip was not sterilized, and the experiment was conducted on a microscope stage. The images were taken at different time points during flow test. Scale bar:  250 µm.

### Supplementary Figure 6

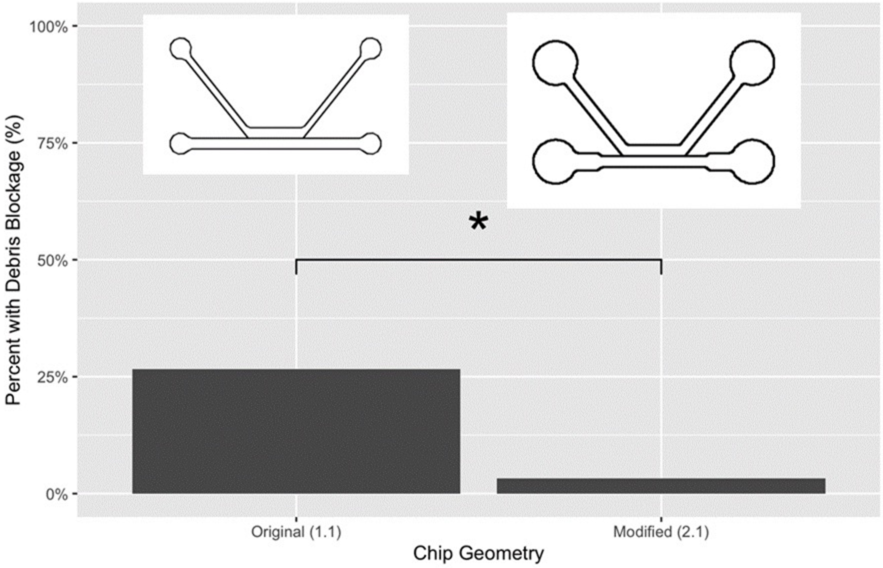


**Supplementary Figure 6. Effect of different channel geometries on the occurrence of channel blockage.** Bar plot comparing the percentage of chips with debris between chip 1.1 and chip 2.1.

### Supplementary Figure 7

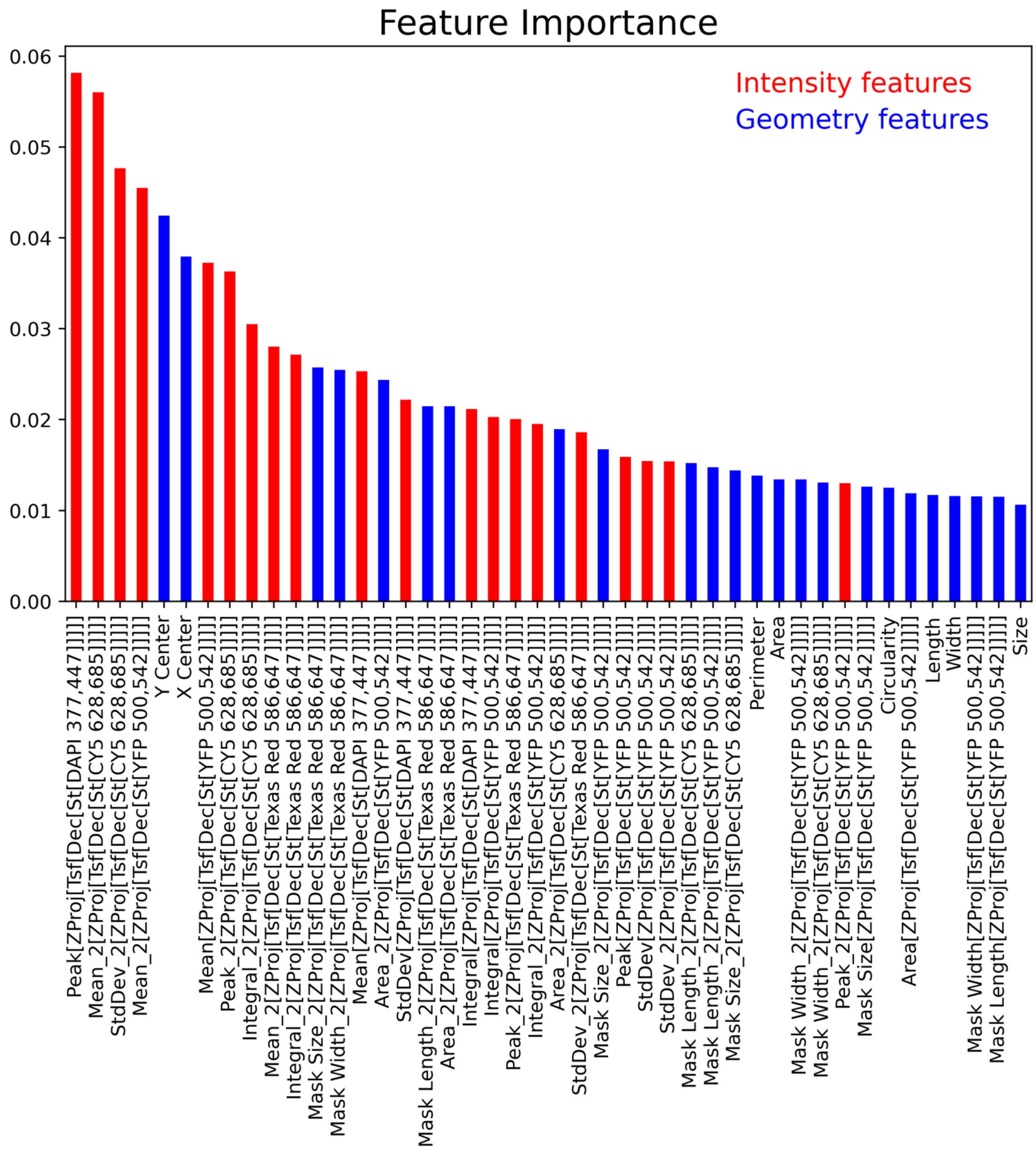


**Supplementary Figure 7. Morphological analysis of CellPainting images.** Bar graph showing feature importance ranking color-coded by either intensity-based features or geometry features. Note that the top-ranking features are intensity features.

| Staining Reagents | Cellular Compartment | Vendor Cat. No. | Imaging Channel | Ex (nm) | Em(nm) |
|---|---|---|---|---|---|
| MitoTracker Deep Red | Mitochondria | Invitrogen #M22426 | Cy5 | 623 | 628/685 |
| Phalloidin/ Alexa Fluor 568 | Cell Membrane | Invitrogen #A12380 | TxRed | 590 | 586/647 |
| Wheat-Germ Agglutinin/ Alexa Fluor 555 | F-Actin Cytoskeleton | Invitrogen, #W32464 | TRITC | 554 | 556/600 |
| SYTO14 Green Fluorescent Nucleic Acid Dye | Nucleoli, Cytoplasmic RNA | Invitrogen, #S7576 | YFP | 505 | 500/542 |
| Concanavalin A/ Alexa Fluor 488 | Endoplasmic Reticulum | Invitrogen #C11252 | GFP | 465 | 469/525 |
| Hoechst 33342 | Nucleus-dsDNA Selective | Thermo Fisher Scientific #62249 | DAPI | 365 | 377/447 |

**Supplementary Table 1. Staining reagents and imaging settings for CellPainting.**

| Staining Solution Type | Component Dye and Concentration | Solvent |
|---|---|---|
| **Live-Cell** | MitoTracker Deep Red (500 nM)<br>Wheat-Germ Agglutinin/Alexa Fluor 555 (1.5 μg/ml) | 1xPBS |
| **Fixed-Cell** | Phalloidin Alexa Fluor 568 (165 nM)<br>SYTO14 Green Fluorescent Nucleic Acid Dye (3 μM)<br>Concanavalin A/Alexa Fluor 488 (50 μg/ml)<br>Hoechst 33342 (20 μg/ml) | 1xPBS |

**Supplementary Table 2. Conditions for staining reagents.**
